# Development of a liposomal formulation of a TKI: the active loading of erlotinib.HCl

**DOI:** 10.1101/2025.06.06.658400

**Authors:** Cindy Schelker, Victoria Trap, Gerrit Borchard, Patrycja Nowak-Sliwinska

## Abstract

Erlotinib.HCl is a tyrosine kinase inhibitor that is used to treat NSCLC and pancreatic cancer in combination with gemcitabine. It is associated with severe adverse effects. This has led researchers to explore encapsulation strategies to reduce these effects. At the same time, previous studies have investigated diverse forms of nanoparticle-based delivery systems for erlotinib.HCl, active loading into liposomes remains unexplored. In this study, we report the first successful active loading of erlotinib.HCl into liposomes.

A critical step in nanomedicine development is its translation to clinical settings. Current studies typically progress from 2D cell culture to mouse models. The former lacks complexity, while the latter lacks relevant human physiology. We believe there is a gap between the two models. Therefore, we evaluated our formulation using both traditional cell lines and patient-derived organoids (PDOs).

Our findings demonstrate successful active loading of erlotinib.HCl into liposomes. Comparative analysis revealed significant differences in both toxicity profiles and cellular uptake between 2D cell lines and PDOs, highlighting the importance of using complementary model systems in nanomedicine development.

## 1. Introduction

Several generations of targeted therapies have emerged over recent years, particularly tyrosine kinase inhibitors (TKIs), which have transformed the treatment landscape of non-small cell lung cancer (NSCLC).

First-generation epidermal growth factor receptor (EGFR) inhibitors such as gefitinib and erlotinib.HCl were followed by second- and third-generation inhibitors such as afatinib and osimertinib, resistance-associated mutations [4]. Erlotinib.HCl binds to the intracellular domain of the protein kinase EGFR, competing with ATP, reducing the receptor’s phosphorylation, and inhibiting the downstream pathway [5]. Erlotinib.HCl is an FDA-approved drug marketed under the name Tarceva^®^ in 2004 for the treatment of locally advanced or metastatic NSCLC after at least one chemotherapy regimen [5]. In 2005, its indication was expanded to include metastatic NSCLC and metastatic pancreatic cancer in combination with gemcitabine [6]. Treatment of NSCLC with erlotinib.HCl led to initial success against lung cancer, but drug-acquired resistance develops after 10-13 months of treatment [7, 8]. These resistance mechanisms limit the long-term efficacy of targeted therapies and highlight the need for novel therapeutic strategies and combination regimens aimed at overcoming or delaying resistance development. A meta-analysis comparing the efficacy of erlotinib.HCl monotherapy to erlotinib.HCl combined with an antiangiogenic agent (bevacizumab or ramucirumab) concluded that the combination significantly improved the progression-free survival (PFS) with a hazard ratio (HR) of 0,59 with 95% confidence interval (0,51-0,69) compared to monotherapy. Unfortunately, long-term treatment with erlotinib.HCl treatment (over six months) can lead to important side effects such as skin toxicity, severe fatigue, or depression, and represents a major issue for erlotinib.HCl treatment [7, 9].

As a result, various strategies of erlotinib.HCl’s encapsulation have been widely documented in the literature, aiming at bioavailability enhancement, side effect reduction, or enabling the co-delivery with another active pharmaceutical ingredient (API) such as paclitaxel, doxorubicin, and quercetin [10–12]. Both *in vitro* and *in vivo* experiments have used a wide range of nanocarriers such as polymer-based, lipid-based, inorganic, and hybrid particles to load erlotinib.HCl [13].

Drug loading within liposomes can be achieved through passive or active/remote processes. The former refers to the API encapsulation as liposomes are formed. The latter occurs when the API is being loaded posterior to liposome formation. Several FDA-approved liposomal products use active loading, such as Doxil^®,^ with doxorubicin loaded using an ammonium sulfate gradient [14], or Vyxeos^©^ with irinotecan complexed with copper ions [15]. Active loading generally leads to higher drug encapsulation efficiencies and reduces leakage during both storage and after parenteral administration, thus increasing the levels of drug the tumor is exposed to [16, 17]. Erlotinib.HCl is a hydrophobic API with two amines on its quinazoline ring and can become positively charged at pH values below 5, increasing the API solubility in aqueous media. In its protonated form, erlotinib.HCl can complex with sulfate ions from ammonium sulfate, precipitating and remaining trapped in the inner aqueous phase of liposomes. This precipitation is advantageous and can limit drug leakage, making erlotinib.HCl a strong candidate for active loading.

A consistent challenge in nanomedicine formulation development is the high rate of clinical translation failure often due to toxicity and lack of efficacy [18]. Toxicity assessments on non-cancerous cells are frequently overlooked, with studies transitioning from *in vitro* cancer models directly to *in vivo*, without adequate evaluation of non-malignant cells. While two-dimensional (2D) cell cultures offer a cost-effective and rapid screening platform for preclinical toxicity evaluation, it lacks the complexity necessary to accurately represent *in vivo* conditions. Similarly, murine models often fail to capture the diversity of human cellular responses. Three-dimensional (3D) models, such as patient-derived organoids (PDOs), provide a more physiologically relevant system for assessing nanomedicine toxicity and efficacy. These models address several limitations of 2D culturing and animal models and could serve as a valuable intermediate step in preclinical testing [19, 20].

To our knowledge, this is the first study to document erlotinib.HCl actively loaded into liposomes. This methodology could lead to stable drug retention and higher encapsulation efficiency [17, 21]. We developed a liposomal formulation encapsulating erlotinib hydrochloride and evaluated its effects through physicochemical characterization and toxicity *in vitro* assessment. These evaluations were conducted on both non-cancerous and cancerous cells in 2D cell cultures, as well as in patient-derived organoids (PDOs).

## 2. Materials and Methods

Erlotinib.HCl was purchased from LC Labs (Woburn, MA, USA). 1,2-dipalmitoylphosphatidylcholine (DPPC) was purchased from Avanti Research (Birmingham, AL, USA), Cholesterol (CHOL), D-α- tocopheryl polyethylene glycol 1000 succinate (TPGS),, 2-Dipalmitoyl-sn-glycero-3-phosphoglycerol (DPPG), Triton X-100^®^, anhydrous DMSO, phosphate salt (H_2_KO_4_P), ammonium sulfate (AS) and sucrose were all purchased from Sigma Aldrich Chemie GmbH (Schnelldorf, Germany). Methanol, acetonitrile, and ethanol were obtained from Fischer Chemical (Reinach, Switzerland). Citric acid monohydrate was purchased from Hänseler AG (Herisau, Switzerland). Sephadex G-25 Hitrap desalting column was purchased from Cytiva (Rosersberg, Sweden), electronic pipette was from Eppendorf (Hamburg, Germany).

### Physicochemical properties of liposomes

#### 2.1 UPLC-UV quantification

The chromatographic separation of erlotinib.HCl was done using a Waters Acquity System (Milford, MA, USA) equipped with a binary solvent delivery pump, autosampler, sample manager, and a photodiode array (PDA) detector. Separation was carried out on an Acquity UPLC^®^ BEH C18 2.1 x 100 mm, 1.7 µm column. The column was heated to 30°C. The mobile phase was composed of solvent A (phosphate buffer: H_2_KO_4_P 0.05M, pH 6.5 adjusted with NaOH) and solvent B (Methanol: Acetonitrile 70:30). A gradient method was used starting from a mixture of 50% of A and 50% of B to 100% of B in 6 min. From 6 min up to 6.1 min the mobile phase was a mixture of 50% A and 50% B and stayed at this composition for 7 min. The flow rate was 0.3 ml/min and UV detection was achieved at 246 nm. The injection volume was 10 µl. Stock solutions of erlotinib.HCl (10 mg/ml) were prepared by dissolving drugs in DMSO, and were stored at -20°C until further use. Calibration solutions of erlotinib.HCl were prepared under the same conditions as the samples.

An example of a chromatograph of erlotinib.HCl separation with UPLC-UV and an example of a calibration curve of erlotinib.HCl in PBS: DMSO 1:4 v/v are displayed respectively in **Supplementary Figure S10** and **Supplementary Figure S11**.

#### 2.2 Calibration curve

A standard stock solution of erlotinib.HCl in anhydrous DMSO at 10 mg/ml was diluted with increasing concentrations of PBS: DMSO 1:4 v/v at pH 7.4. The final range of concentrations was 0.01 – 0.02 – 0.05 – 0.10 – 0.2 – 1 µg/ml. The calibration curve of erlotinib.HCl was plotted with the area under the peak on the Y axis and erlotinib.HCl concentration on the X axis.

#### 2.3 Drug loading and entrapment efficiency

Erlotinib.HCl presence was assessed either with a microplate reader (BioTek Instruments, Sursee Switzerland) or with a UV absorbance spectrum. Erlotinib.HCl quantification in liposomal formulations was determined using UPLC-UV. 20 µL of formulation was diluted with DMSO to a final volume of 200 µL in a 2-mL glass vial prior to determination by the UPLC-UV.

The encapsulation efficiency (EE%) was calculated using Equation 1.

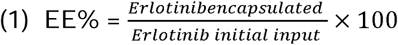

#### 2.4 Liposome preparation

Liposomes were prepared using the ethanol injection method. Two different protocols were followed to search for optimal loading conditions.

For erlotinib.HCl passive loading, 50 µL of erlotinib.HCl (10 mg/ml dissolved in DMSO), DPPC (1.42 mg/ml), DPPG (0.24 mg/ml), Cholesterol (1,63 mg/ml) and TPGS (0.20 mg/ml) were added to ethanol with Kolliphor^®^. The ethanolic mixture was added under stirring at 600 rpm with an electronic pipette to PBS at pH 7.4 and heated at 60°C. The mixture was cooled down on ice. Free erlotinib.HCl, DMSO and ethanol were removed by size exclusion chromatography (SEC) with a Sephadex G-25 column.

For erlotinib.HCl active loading, lipids were dissolved in ethanol with or without Kolliphor^®^. The final composition of lipids in ethanol was DPPC (1.42 mg/ml), DPPG (0.24 mg/ml), Cholesterol (1,63 mg/ml), TPGS (0.20 mg/ml) and Kolliphor^®^ HS-15 (0,04 mg/ml). The ethanol mixture was rapidly injected by an electronic pipette in ammonium sulfate buffer at pH 3 heated to 60°C. Freshly prepared liposomes were put on ice to cool down. Then, external ammonium sulfate was exchanged for citrate buffer at 0,1 M/sucrose 10% by SEC. Liposome suspensions were then returned to heat at 60°C in balloons and 50 µL erlotinib.HCl (10 mg/ml) was added. The mixture was left under stirring for one hour and then cooled on ice. A second SEC was performed to exchange citrate/sucrose buffer to PBS and remove free erlotinib.HCl.

Blank liposomes were identically prepared without the addition of the API.

Fluorescently labeled liposomes were prepared following the active loading protocol with the replacement of 1 mol% of Cholesterol with NBD-Cholesterol, a fluorescent cholesterol molecule (Ex/Em 469/537 nm).

#### 2.5 Hydrodynamic diameter, polydispersity index (PDI) and surface charge

Mean hydrodynamic size (Z-average), PDI and surface charge (Zeta-potential) were determined by dynamic light scattering using a Zetasizer (NanoZS), Malvern Panalytical, Malvern, UK) in batch mode. Samples were measured at 25°C in polystyrene disposable cuvettes. Measurement angle was at 173°C, the refractive index was set at 1.345, and absorption at 0.010. The laser attenuator was adjusted automatically. Measurements were done in triplicates. The equilibration time between samples was 120 seconds. Size and PDI values were measured after dilution at a ratio of 1:20 with PBS 0.9% filtered through filters of a 0.22 µm pore size. Zeta potential was measured after dilution at a ratio of 1:100 with Milli-Q^®^ water. Results are presented as mean ± SD. Data were collected using Zetasizer software v7.13.

#### 2.6 *In vitro* drug release

Erlotinib.HCl solubility was first assessed in the release medium (PBS/Tween 80 0.1% at 37°C). Erlotinib.HCl was dissolved in excess in release media and after 72 hours, drug content was quantified by UPLC. Erlotinib.HCl release from liposomes was then assessed. First, the liposomal suspensions were concentrated: 4 mL of liposome suspension was added to centrifugal filters (Sigma-Aldrich, 6 mL, MWCO: 10 kDa). Centrifugation was at 4,000 rpm for 80 min at 4°C. The concentration of erlotinib.HCl was then assessed with UPLC-UV to determine the starting concentration. 0.5 mL of suspension was put in a dialysis cassette with MWCO at 10 KDa (Slide-a-lyzer™, Thermo Fisher Scientific, Carlsbad, CA, USA). The release medium was PBS/Tween 80 0.1% at pH 7 with a volume of 50 mL. Aliquots of 200 µL were taken at each timepoint.

#### 2.7 Stability studies

Drug retention, sizes, PDI and zeta-potential were followed as a function of time. Liposomal suspensions were stored at 4°C.

#### 2.8 Cryogenic transmission electron microscopy (Cryo-TEM)

Liposomal morphology was evaluated by Cryo-TEM. Perforated carbon grids with 200-mesh density (Quantifoil, Hatfield, PA, USA) were glow-discharged using an EMS GlowQube instrument. Liposomes were diluted 20-fold with PBS and deposited on the grids. Grids containing the samples were vitrified by immersion in liquid ethane using a Leica EM GP2-Automatic Plunge Freezer (Wetzlar, Germany) and imaged with a Talos Arctica (200 KeV, FEG, Thermo Fisher Scientific, Carlsbad, CA, USA). Images were analyzed with Fiji software (Image J, 1.54f).

### Biological characterization of liposomes

#### 2.9 Cell culture and PDO generation

Human embryonic kidney 293T cells (HEK-293T), and human metastatic RCC cell line (786-O) were purchased from American Type Culture Collection (ATTC, Teddington, UK). They were cultured in RPMI 1640 medium supplemented with 10% (v/v) of heat-inactivated fetal bovine serum (FBS) and 1% of penicillin/streptomycin (Gibco, Thermo Fisher Scientific, Carlsbad, CA, USA).

The patient-derived material was obtained via the protocol approved by Swiss ethics committees on research involving humans (2017-00364) with signed patient consent. Protocol to generate PDOs from human kidneys and RCC tissues was performed as published previously [22].

#### 2.10 Mitochondrial activity (WST-1)

HEK-293T and 786-O cells were seeded at an initial density of 5.10^5^ cells/well in 96-well plates and allowed to attach for 24h. Cells were treated with either NaCl 0.9% in cell culture medium as control for “blank liposomes” and “liposomal formulation of tubacin” (CTRL) or NaCl 0,9% in cell culture medium with 0.1% DMSO as a control for the “free tubacin” condition. Blank liposomes, free tubacin, or liposomal tubacin and incubated for 24h. After the incubation, the medium was aspirated. 100 µL of WST-1 reagent (4-[3-(4-iodophenyl)-2-(4-nitrophenyl)-2H-5-tetrazolio]-1,3-benzene disulfonate, Roche, Basel, Switzerland) diluted 1/10 with medium were added to each well with the cells and incubated for 30 min in the cell incubator at 37°C in a humidified atmosphere with 5% CO_2_. UV absorbance was measured at λ = 450 nm using a microplate reader (BioTek Instruments, Sursee, Switzerland). CTLR were used as a positive reference corresponding to 100 % viability. Mitochondrial activity is displayed as a mean ± standard deviation (SD) of three biological replicates (N=3). Equation 1 was used to calculate the mitochondrial activity of every sample:

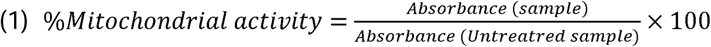

#### 2.11 CellTiter-Glo^®^

1000 cells/well were seeded in 96-well U-bottom low attachment plates (650970, Greiner). The cell culture medium of the PDOs was Renal epithelial cell growth medium 2 (Promocell) for the kidney organoids and DMEM/F12 supplemented with Primocin, Stempro (Gibco), Noggin, Wnt3a, hEGF, hFGF2, hFGF10, PDGFbb, R-spondin1, R-spondin3. Both media were supplemented with 2.5% Matrigel^TM^ (354254, Corning) before addition to the plate. Plates were centrifuged for 5 minutes at 400 x g at 4°C immediately after seeding. After 4 days of incubation at 37°C, 70% of humidity and 5% CO_2_, PDOs were treated. Cell metabolic activity assay was measured with CellTiter-Glo^®^ assay (Promega, Madison, USA). 50 µl of CellTiter-Glo^®^ were added to each well and incubated at room temperature in the dark for 30 minutes. Then 100 µl of samples were transferred to a black, flat-bottom 96-well plate and luminescence was detected using the Biotek Cytation 5 (BioTek Instruments, Sursee, Switzerland) using corresponding software at default settings. Equation 2 was used to calculate the ATP levels of every sample:

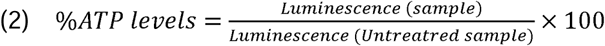

### Cellular uptake

Cells were seeded in µ-Slide 8 Well Ibidi plates (Ibidi GmbH, Baar, Switzerland) at a density of 2,5.10^4^ cells per well and incubated at 37°C, 5% CO_2_ in complete growth medium until 70% confluence. Fluorescent liposomes were added to each well and incubated for 5 min to 3 hours. Following incubation, cells were washed twice with PBS and fixed in 4% paraformaldehyde for 10 min at ambient temperature.

Following fixation cells were washed with PBS twice. Cells were counterstained with Hoechst 33342 (Thermofisher, Basel, Switzerland). A drop of VECTASHIELD^®^ mounting medium was added to reduce photobleaching. Finally, visualization of cellular uptake was studied using the confocal laser scanning microscopy (CLSM) technique. The slides were examined on a Nikon A1r Spectral point scanning confocal microscope with either a 10x (for PDOs) or 60x (for cells cultured in 2D) oil immersion objective for cells in a 2D model. Images were analyzed with Fiji software (Image J, 1.54f).

#### 2.12 Statistical analysis

Data analysis was performed using GraphPad Prism software version 10.2.3. A two-way ANOVA was performed, and the data between groups was compared using Tukey’s multiple comparisons test. All *p* values < 0.05 were considered statistically significant.

## 3. Results

### 3.1 Preliminary results

#### Liposomal erlotinib.HCl loading

Erlotinib.HCl is a hydrophobic small molecule with a log P of 3.2 (**Figure 1A**), making it a suitable candidate for liposomal encapsulation. Liposomes were synthesized via the ethanol injection method, and both passive (**Figure 1B)** and active loading (**Figure 1C**) strategies were investigated. The formulation comprised DPPC:DPPG:CHOL:TPGS:KOLL in a 47:8:20:3:1 molar ratio.

**Figure 1.**
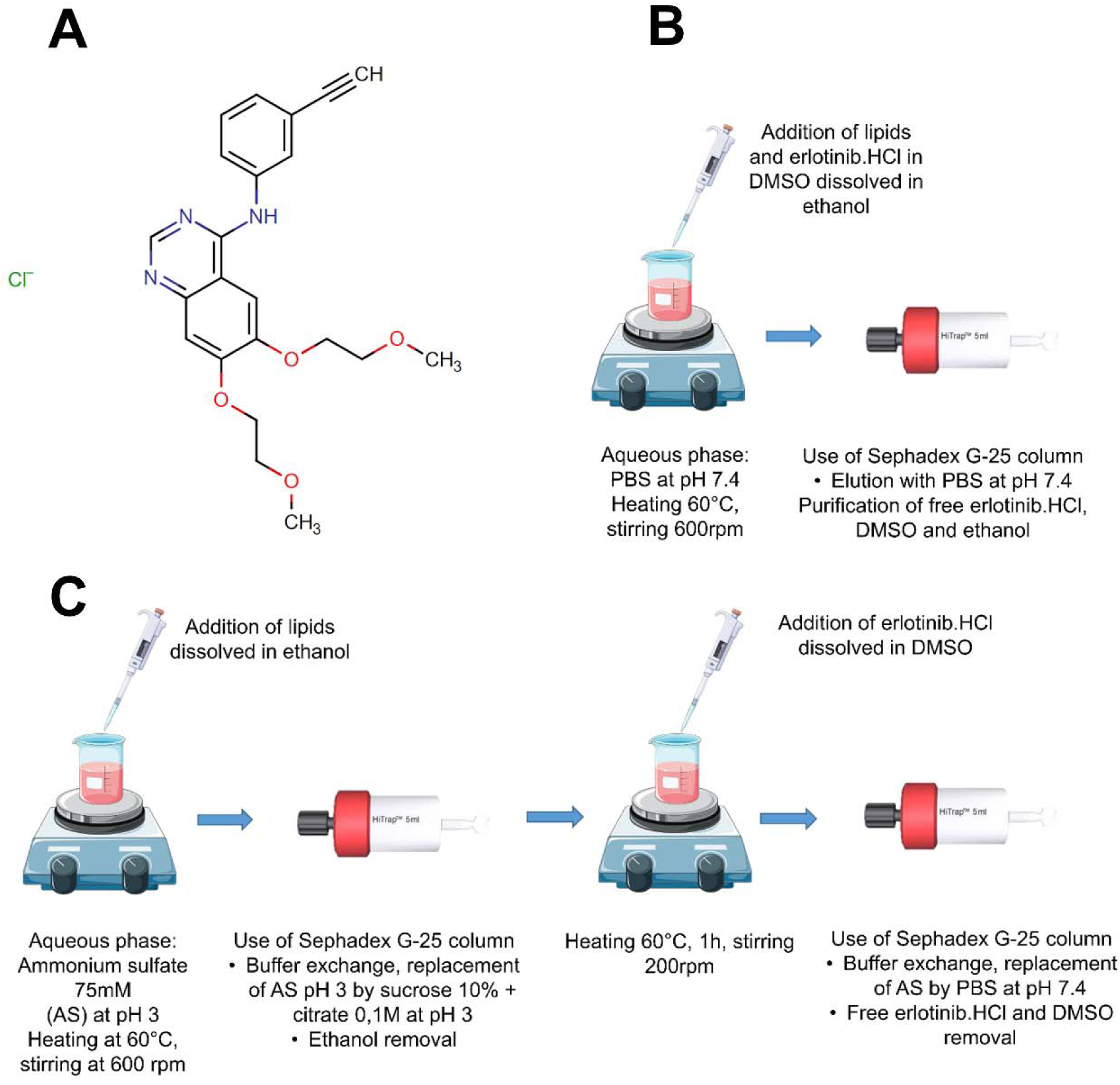
Erlotinib hydrochloride salt structure from Pubchem compound database (http://pubchem.ncbi.nlm.nih.gov/): erlotinib (#176870) (**A**). Schematic representation of passive loading method for the preparation of erlotinib.HCl-loaded liposomes (**B**). Schematic representation of active loading method for the preparation of erlotinib.HCl-loaded liposomes (**C**).

The absorbance spectrum of liposomes exhibited peaks at 280 and 350 nm. As confirmed by UPLC-UV analysis, the 350 nm peak confirms the presence of erlotinib.HCl (results not shown). This peak is absent in conditions such as passive loading or active loading without cholesterol or Kolliphor^®^, marking the lack of erlotinib.HCl encapsulation (**Figure 2A**). Following excitation at 350 nm (**Figure 2B**) an emission peak at 380 nm was detected in both drug-loaded and blank liposomes, suggesting that this signal was not attributable to erlotinib.HCl. Interestingly, actively loaded liposomes containing both cholesterol and Kolliphor^®^ spectra had two distinct peaks: one at 446 nm and one at 480 nm.

**Figure 2.**
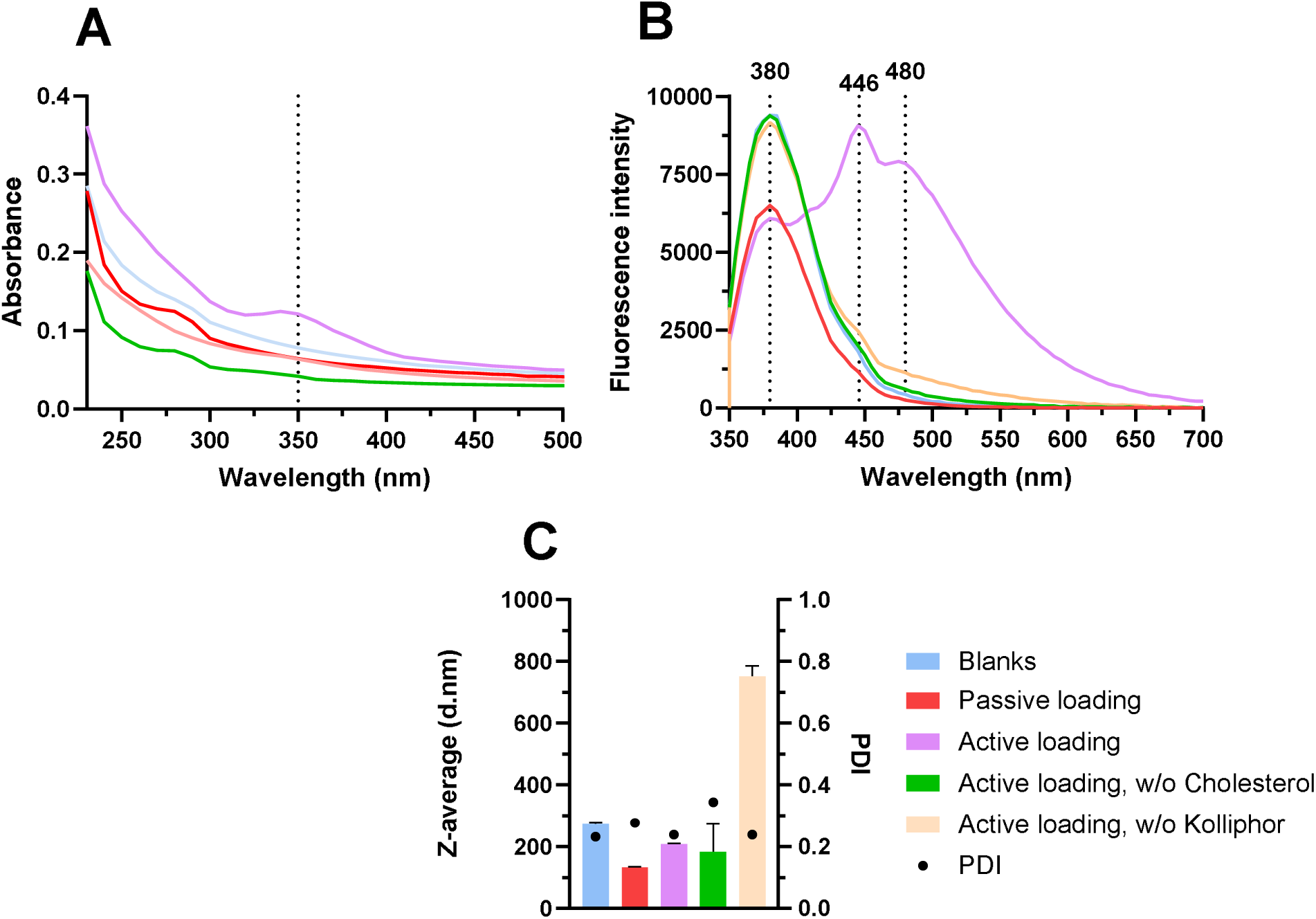
Absorbance (**A**) and emission spectra (**B**) and Z-average/PDI of erlotinib.HCl-loaded liposomes prepared with varying conditions (**C**), N=1.

The two peaks were absent in blank liposome preparations, confirming their association with erlotinib.HCl encapsulation. No emission was detected under other conditions. The z-average obtained ranged between 100 and 200 nm across all formulations, except in the absence of Kolliphor^®^ where particles increased to 800 nm (**Figure 2C**). Following these preliminary findings, liposomes were subsequently prepared via active loading in the presence of both Kolliphor^®^ and cholesterol.

### 3.2. Physicochemical characterization of erlotinib.HCl-loaded liposomes

The stability of sizes was evaluated during storage at 4°C. Sizes measured by DLS by intensity ranged between 200 and 300 nm, with PDIs remaining below 0.2 from the day of manufacturing (day 0) to day 7 (see **Figure 3A**). Size measurements by number revealed a population with diameters around 100 nm over the same period and storage conditions (see **Figure 3B**). This dual representation, size by intensity and size by number, was further confirmed by the Zetasizer software’s detailed report (see **Figure 3C** and **3D**), which showed that 86 to 90% of the particles, measuring 95 nm, remained stable for at least 7 days at 4°C.

**Figure 3.**
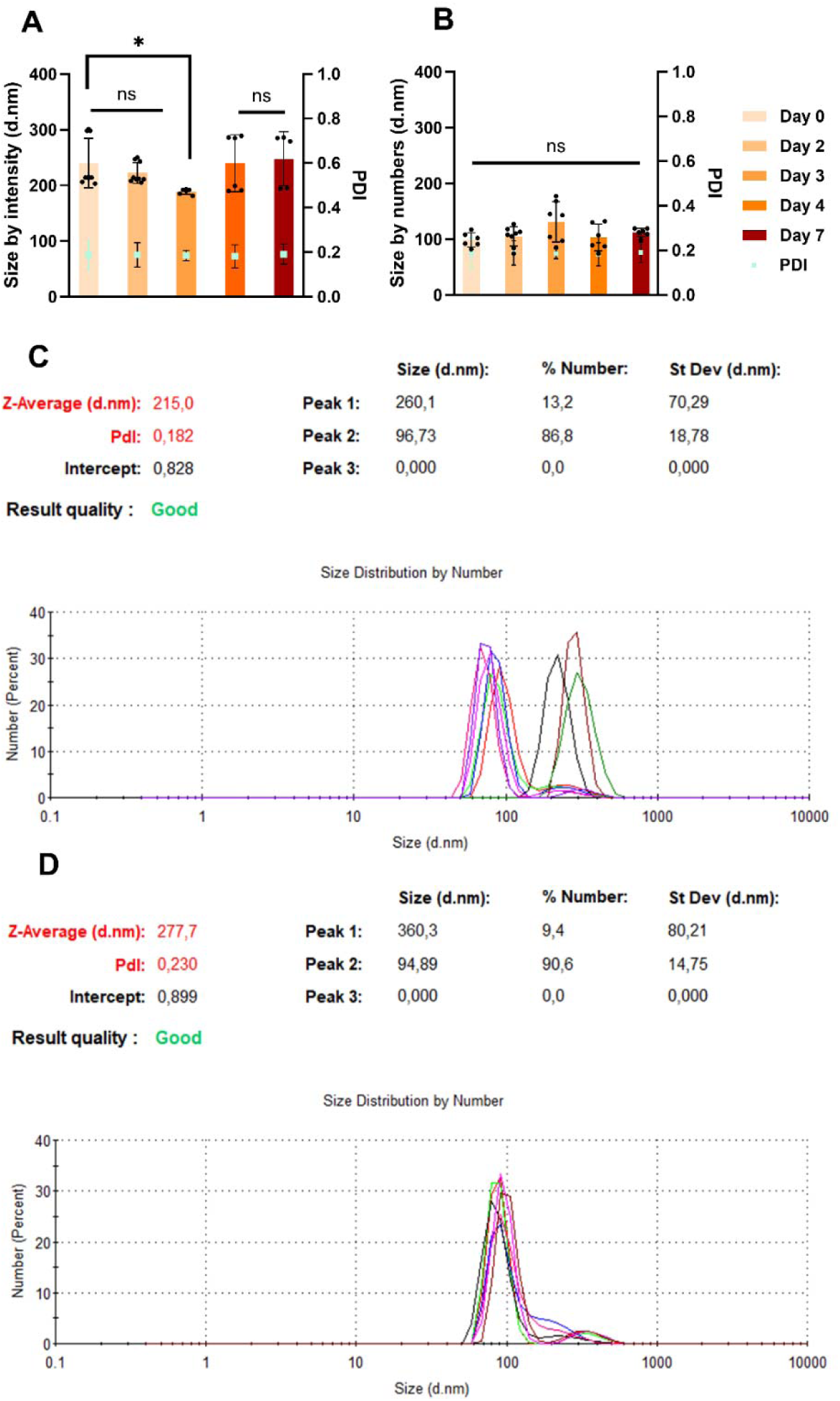
Size characterization of DPPG:DPPC:CHOL:TPGS:KOLL liposomes (**A**) Erlotinib.HCl-loaded liposomes sizes by intensity. N=3 ± SD (**B**) Erlotinib.HCl-loaded liposome sizes by number N=3 ± SD. Characterization of erlotinib.HCl-loaded liposomes. Size by number the day of the synthesis, day 0 (**C**), and after 7 days of 4°C storage (**D**).

Erlotinib.HCl’s spectral properties were analyzed using a spectrophotometer to rapidly verify drug presence and confirm preliminary findings. Analysis of loaded liposomes revealed the characteristic emission peak of erlotinib.HCl at 350 nm (see **Figure 4A).** Fluorescence emission monitoring showed two emission peaks at 446 and 480 nm, confirming encapsulation in liposomes (see **Figure 4B).** Encapsulation efficiency determined using UPLC-UV by plotting the area under the curve against known standard concentrations from a calibration curve (see **Figure 4C**). The formulation had a stable negative surface charge of -40 mV over 7 days at 4°C (see **Figure 4D**). (Encapsulation efficiency was 35%, and drug release during storage was assessed. A 15% loss of encapsulated erlotinib.HCl was observed during the first 48 hours, with no additional loss recorded for at least 168 hours (7 days) at 4°C (see **Figure 4E**). To evaluate the release after systemic administration, drug release kinetics were assessed at 37°C. Erlotinib.HCl was soluble in the release medium up to 0.028 mg/ml with a maximum concentration of 0.00058 mg/ml in the medium to maintain sink conditions. Finally, the drug release profile at 37°C displayed a burst release of erlotinib.HCl from liposomes within the first 4 hours, followed by a stable phase with no further release for 120 hours (5 days). The final erlotinib.HCl content was fully released after 168 hours (7 days) (see **Figure 4F**). Comparatively, free erlotinib.HCl was completely released from the dialysis cassette within 5 hours.

**Figure 4.**
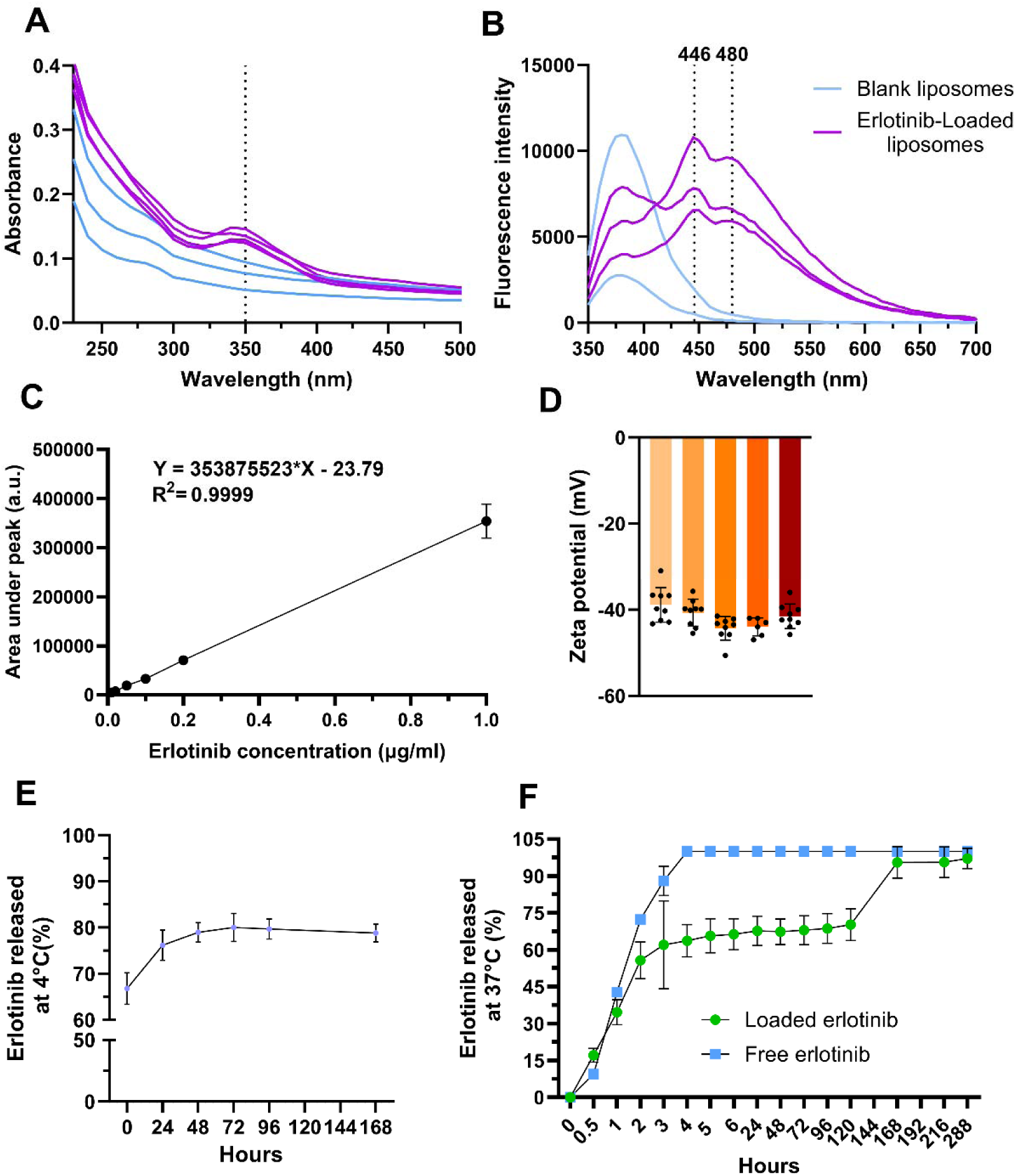
Physico-chemical characterization of DPPG:DPPC:CHOL:TPGS:KOLL liposomes. Absorbance and emission spectra of both blank liposomes and erlotinib.HCl–loaded liposomes (**A)** Absorbance spectra. (**B**) Emission spectra, λ(excitation) = 350 nm. N=3. (**C**) Calibration curve obtained with UPLC-UV for the quantification of erlotinib.HCl. N=1, error bar corresponds to SD (**D**) Erlotinib.HCl-loaded liposomes ζ-potential for 7 days stored at 4°C. N=3, error bar corresponds to SD. (**E**) Erlotinib.HCl encapsulation stability of stored liposomes at 4°C. N=2, error bar corresponds to SD. (**F**) *In vitro*, using the dialysis cassette method, the drug release profile of free erlotinib.HCl and loaded erlotinib.HCl in PBS/Tween 80 0.1% at 37°C. N=2, error bar corresponds to SD.

Cryo-TEM performed on blank liposomes (**Figure 5A**) revealed the presence of particles containing several bilayers. Size distribution was polydisperse, with some liposomes exhibiting sizes approximately double or triple those of others. The same observations were made for loaded liposomes (**Figure 5B),** which additionally displayed electron-dense, tube-shaped structures.

**Figure 5.**
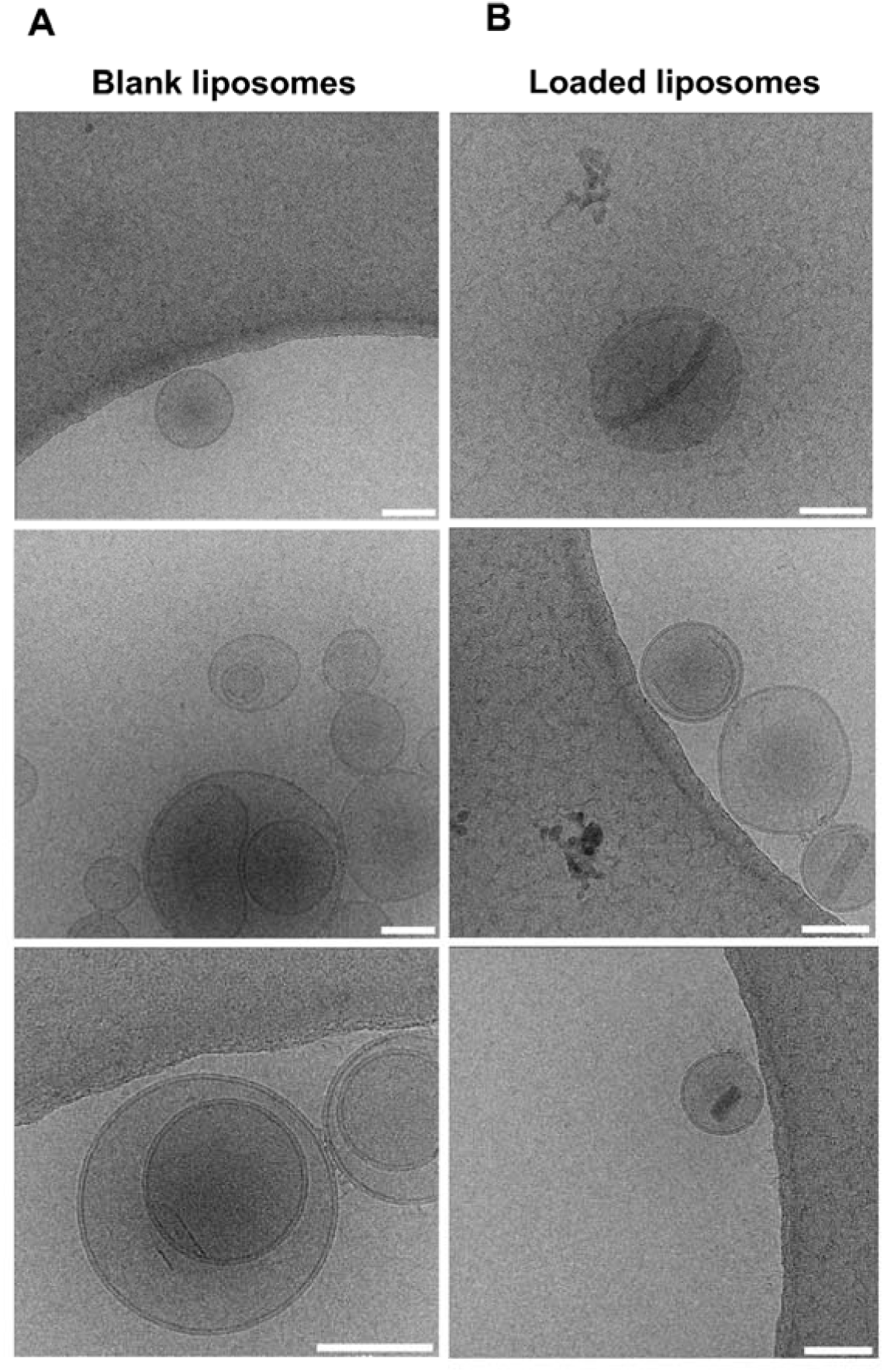
Cryo-TEM images of blank liposomes (**A**) and erlotinib.HCl-loaded liposomes (**B**). Scale bar = 100 nm.

### 3.3. Efficacy and safety assessment of erlotinib.HCl-loaded liposomes on 2D and 3D cell cultures

The toxicity of erlotinib.HCl-loaded liposomes was evaluated using both two-dimensional (2D) cell cultures and three-dimensional (3D) patient-derived organoid (PDOs) models. Assessments were performed on both cancerous and non-malignant systems to determine potential selective cytotoxicity. The 2D cell lines used were 786-O (ccRCC) and HEK-293T (non-malignant, embryonic origin).

In 2D culture, no difference in mitochondrial activity was observed between 786-O and HEK-293T cells after 24 hours of treatment (**Figure 6A**). However, after 72 hours, mitochondrial activity in HEK-293T cells decreased to 40% following treatment with erlotinib.HCl-loaded liposomes, whereas 786-O cells exhibited no reduction (**Figure 6B**). In contrast, free erlotinib.HCl did not result in significant differences in mitochondrial activity between healthy and cancerous cell lines at either 24 or 72 hours (**Figure 6A** and **6B**).

**Figure 6.**
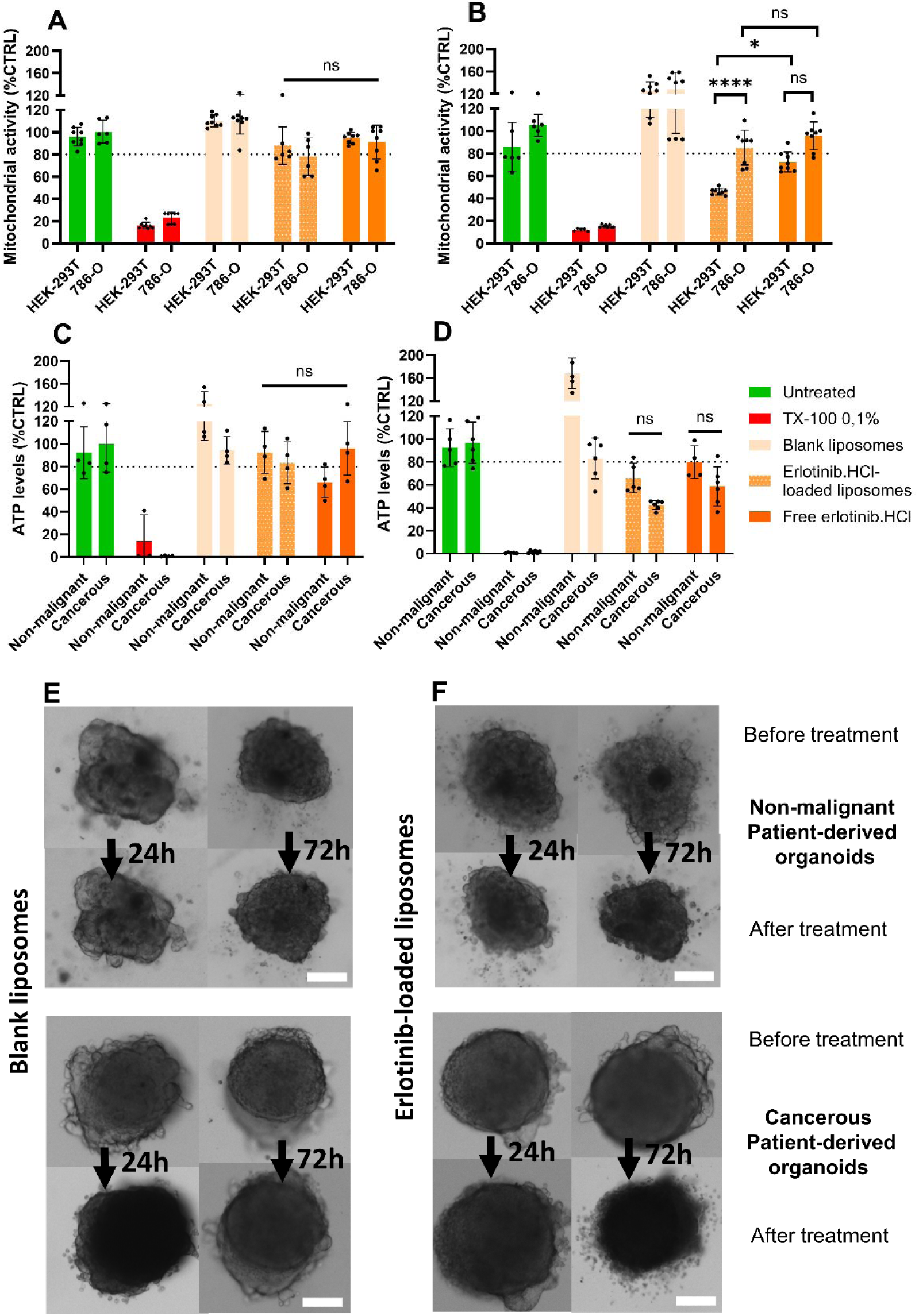
Mitochondrial activities of 2D cell lines HEK-293T and 786-O after 24 hours of treatment (**A**) and 72 hours of treatment (**B**). Data are represented as mean ± SD (N=3). ATP levels of patient-derived organoids (PDOs) from non-malignant and cancerous tissues after 24 hours of treatment (**C**) and 72 hours of treatment (**D**). Data are represented as mean ± SD (N=2). Data were normalized to untreated CTRL. Bright-field images from both non-malignant and cancerous PDOs before and after treatment duration of 24 hours and 72 hours with Blank liposomes (**E**) and erlotinib.HCl-loaded liposomes (**F**). Scale bar = 200 µm. * p ≤ 0.05, **** p ≤ 0,0001, ns stands for non-significance.

In PDOs, ATP levels remained around 80 % after 24 hours of treatment with erlotinib.HCl-loaded liposomes, consistent with observations in 2D cell cultures (**Figure 6C**). However, after 72 hours, no significant difference in ATP levels was observed between cancerous and non-cancerous PDOs, unlike findings in 2D models (**Figure 6D).**

In both 2D and 3D models, blank liposomes demonstrated no toxicity towards non-cancerous or cancerous systems, confirming their inert nature.

The microscopic morphology of organoids was examined following treatment with either blank liposomes or erlotinib.HCl-loaded liposomes at 24 and 72 hours. Blank liposomes did not cause cell detachment in healthy or non-malignant organoids (**Figure 6E**). In contrast, small cell aggregates were observed around organoids treated with erlotinib.HCl-loaded liposomes, particularly after 72 hours (**Figure 6F**), suggesting at least some toxic effect of the drug-loaded liposomes.

### 3.4. Liposomes and drug cellular uptake assessment on 2D and 3D cell cultures

Cellular uptake of liposomes was assessed on both 2D and 3D models. Liposomes were fluorescently labeled with NBD-cholesterol. After incubation with liposomes, cells were washed and fixed.

For HEK-293T (non-malignant) cells, the minimal fluorescence signal was detected from 5 min to 120 min of incubation (**Figure 7A**). In contrast, in 786-O cells, the signal appeared after 45 min and peaked at 120 min (**Figure 7B**). This result indicates that liposomes have a better ability to be taken by 786-O cells, compared to HEK-293T cells. Erlotinib.HCl concentration in the supernatant remained stable during the first hour but decreased between 1 and 3 hours (**Figure 8**).

**Figure 7.**
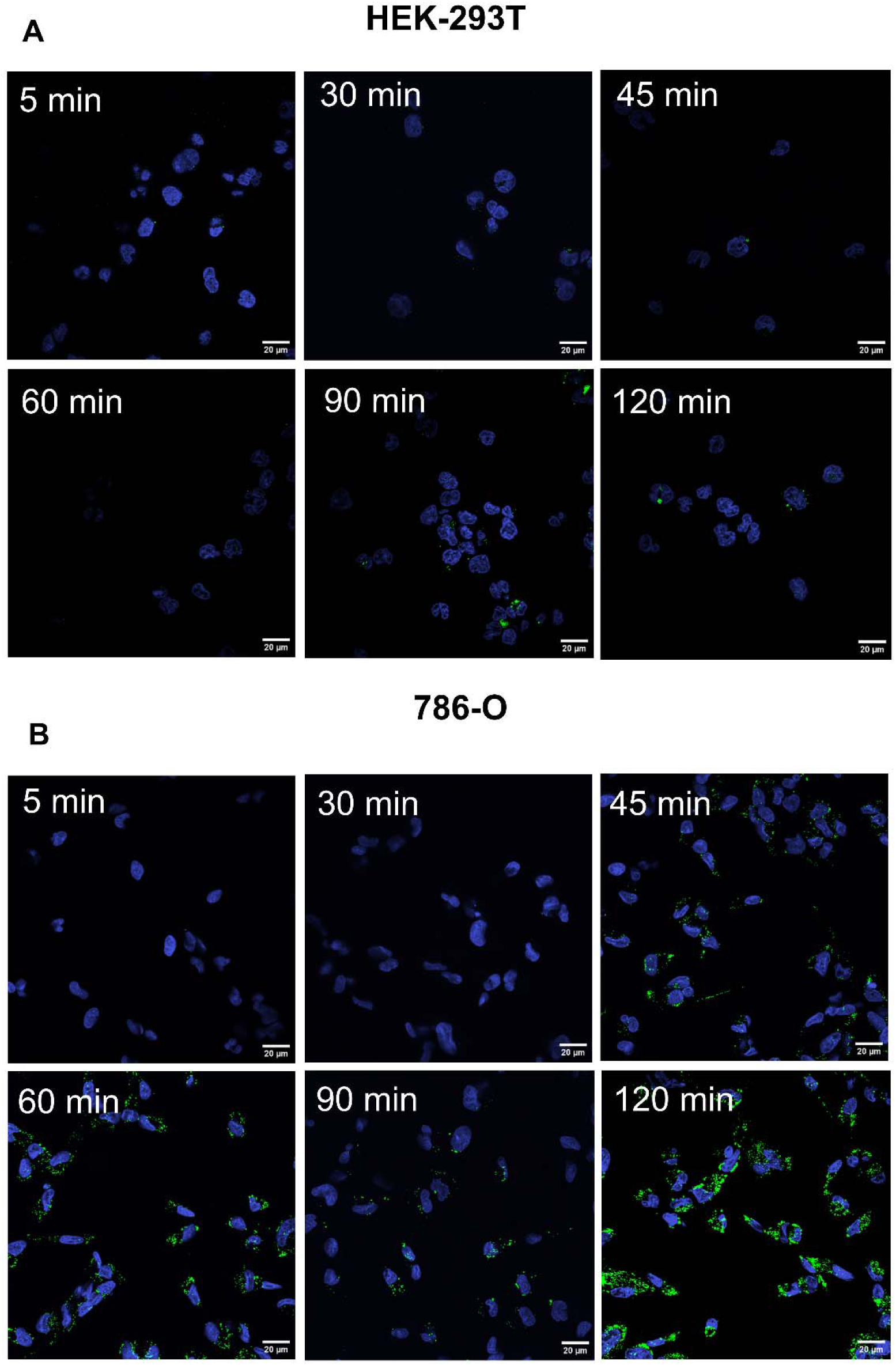
Representative confocal microscopy images of fixed 2D cells after various incubation times with fluorescently labeled liposomes (green). Non-cancerous (HEK-293T) (**A**) and cancerous (786-O) cells (**B**). Scale bar = 20 µm.

**Figure 8.**
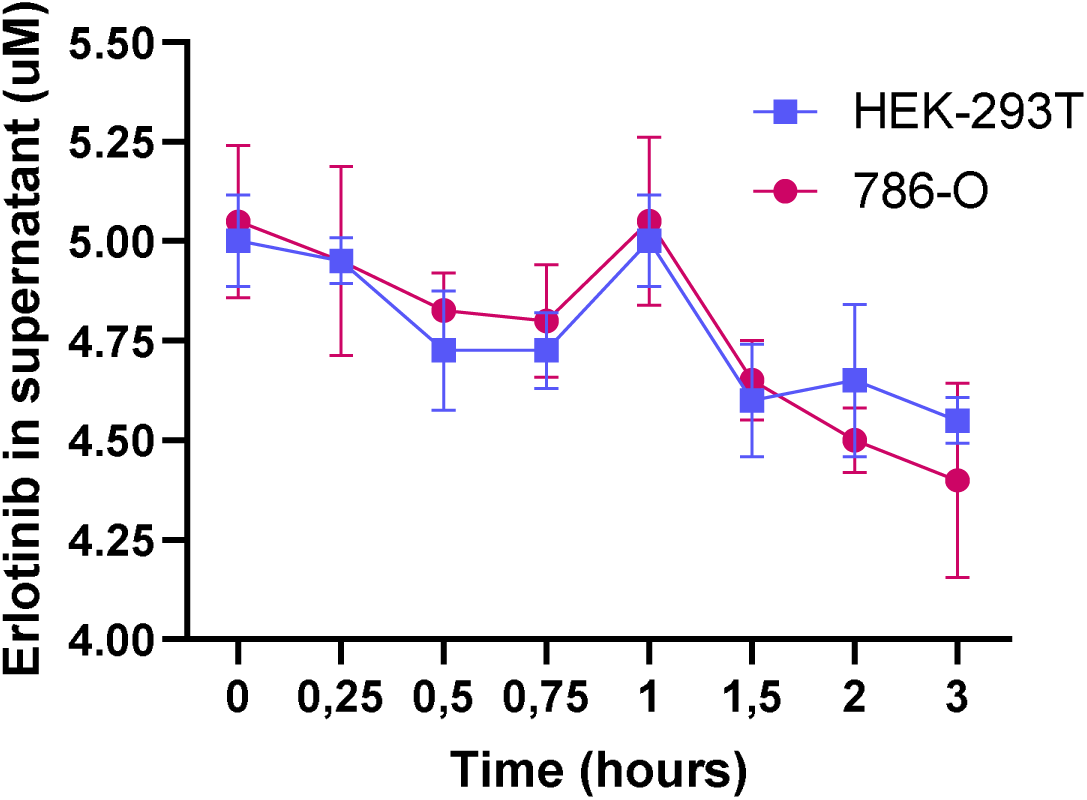
Erlotinib.HCl concentration in the supernatant in 2D cell culture after various incubation timepoints with erlotinib.HCl-loaded liposomes on non-cancerous cells (HEK-293T) and cancerous cells (786-O). Data are represented as mean ± SD (N=2).

Cellular uptake studies using fluorescently labeled liposomes were also performed on PDOs, with confocal imaging focused on the central plane of each organoid (**Figure 9A**). In non-cancerous organoids, erlotinib.HCl-loaded liposomes were internalized within 15 min of incubation (**Figure 9B**). Similarly, in PDOs derived from cancerous tissue, cellular uptake occurred within 15 min, with a consistently higher staining signal at all time points compared to non-cancerous organoids (**Figure 9C**). Liposome uptake was maintained for at least 180 minutes in both cancerous and non-malignant organoids.

**Figure 9.**
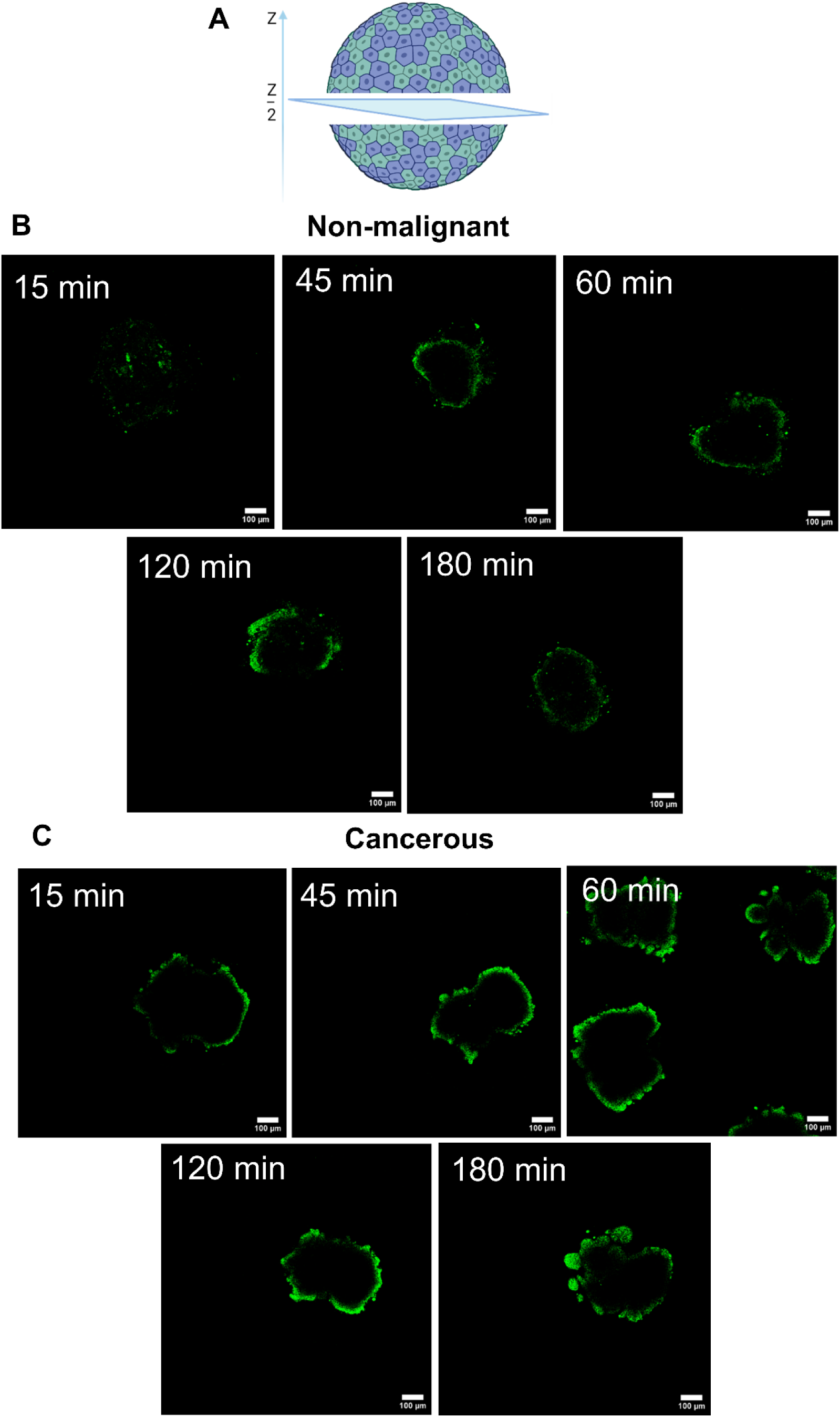
Representative confocal microscopy images of fixed PDOs after different timing of incubation times with fluorescently labeled liposomes (green). Schematic representation of the Z-stack where the confocal plane was placed (**A**). Non-cancerous organoids (**B**) and cancerous organoids (**C**). Scale bar = 100 µm.

## 4. Discussion

Passive loading of erlotinib.HCl has been previously documented in several studies using the thin-film hydration method [23–27]. This methodology consists of dissolving erlotinib.HCl with lipids in ethanol, followed by ethanol evaporation. During this step, the lipid mixture surrounding erlotinib.HCl creates an environment favorable for its solubilization and encapsulation.

In this study, the ethanol injection method (EIM) was used for liposome preparation. EIM was chosen for its potential scalability with microfluidics, as both techniques rely on nanoprecipitation to form liposomes [28]. Despite being a simple and scalable technique [29], EIM has relatively low encapsulation efficiency [30]. This may be explained by the absence of an evaporation step that otherwise enhances hydrophobic drugs, e.g., erlotinib.HCl, interactions with lipids, improving solubilization.

In contrast to passive loading, active loading typically results in higher drug encapsulation and retention [17]. This method requires different medium conditions between the internal phase of liposomes and their exterior. Depending on the physicochemical characteristics of the API, the use of pH gradients or metal complexes can be employed. As illustrated in **Figure 1C**, the active loading process involves two distinct steps: the initial formation of liposomes, followed by external buffer exchange from ammonium sulfate to sucrose 10% + citrate buffer 0,1M pH 3 by size-exclusion chromatography (SEC). Then erlotinib.HCl was added and loaded for 1 hour at 45°C. A second SEC was necessary to remove DMSO, free erlotinib.HCl and exchange the external buffer to PBS.

Erlotinib.HCl’s characteristic UV absorption peak at 350 nm was absent in samples lacking cholesterol and Kolliphor^®^, suggesting that both components are essential for effective drug loading. Kolliphor^®^ HS-15 is a non-ionic surfactant widely used in pharmaceutical formulations to enhance small-molecule solubilization [31]. As an FDA-approved surfactant for parenteral APIs, it consists of a lipophilic part with polyglycol mono- and di-esters of 12-hydroxystearic acid and a hydrophilic part with polyethylene glycol (PEG) chains [32]. During liposome formation, Kolliphor^®^ HS-15 may be incorporated into the bilayer, disrupting phospholipid packing and thereby enhancing drug encapsulation either within the bilayer or the aqueous core. Conversely, cholesterol is known to increase the packing of phospholipids, reducing the bilayer’s permeability to solutes and reducing particle aggregation, making it a critical component in liposomal formulations [33]. It is hypothesized that cholesterol contributes to solute retention and helps maintain the pH gradient, promoting erlotinib.HCl encapsulation [34]. While Kolliphor^®^ HS-15 and cholesterol exert opposing effects on bilayer permeability and stiffness, their combined action facilitates both erlotinib.HCl entry into liposomes and its subsequent retention.

Erlotinib.HCl exhibits negligible intrinsic fluorescence in aqueous media, and this characteristic is not influenced by its protonation state. However, its fluorescence becomes moderate in apolar environments such as n-hexane [35], acetonitrile, and chloroform [36]. Given erlotinib.HCl’s fluorescence is environment-dependent and absent in water; the presence of the 446 and 480 nm peaks may indicate the presence of erlotinib.HCl within an apolar environment, such as the liposome bilayer. These two peaks likely reflect different interactions between erlotinib.HCl and surrounding compounds within that hydrophobic environment.

Regarding particle sizes, Kolliphor^®^ HS-15 presence is reported to reduce particle size. As a surfactant, it lowers the interfacial tension between hydrophobic compounds and water during the liposome formation, thereby promoting smaller particle formation [37]. This is consistent with PDI values observed (∼0,2), which generally indicate a monodisperse size distribution [38]. However, this conclusion is contradicted by Cryo-TEM images revealing a polydisperse size distribution. DLS measurements showed sizes ranging from 200-300 nm when reported by intensity. In contrast, when reported by numbers, 86-90 % of the particles were slightly below 100 nm. Larger particles detected by intensity-based measurements may correspond to aggregates formed during synthesis. The discrepancy arises from the fact that DLS estimates particle size based on light scattering intensity, which is proportional to the sixth power of particle diameter (Rayleigh scattering). This results in larger particles disproportionately influencing intensity-based measurements [39]. While size distributions are commonly reported by intensity in the literature, presenting number-based data provides complementary insight into the actual population composition.

For intravenous (i.v.) administration, most recommendations in the literature advise a nanoparticle size between 100-150 nm to ensure prolonged blood circulation and increase the enhanced retention and permeation (EPR) effect [28].

The zeta potential of the particles was measured at approximately -40 mV, indicating a highly negative surface charge. This charge is primarily attributed to DPPG, a negatively charged lipid, whereas DPPC is a neutral lipid [40]. Zeta potential reflects the charge at the slipping plane, where ions are associated with the liposomes and move collectively as a single entity [41]. Surface charge plays an important role in influencing the biological behavior of liposomes.

Cationic liposomes tend to accumulate more efficiently in tumor vasculature and tissues compared to anionic or neutral liposomes due to electrostatic interactions with the negatively charged glycocalyx lining endothelial cells of microvessels [42]. Nevertheless, while anionic liposomes are also reported to extravasate to tumor tissues, cationic liposomes preferentially accumulate in tumor vessels [43]. However, cationic liposomes also exhibit inherent toxicity, primarily due to their ability to generate reactive oxygen species and activate the complement system. This activation triggers rapid clearance by macrophages, thereby reducing circulation time and limiting efficient tumor delivery [44]. Additionally, the literature suggests the incorporation of PEG-lipid concentrations as low as 1.6 mol% can significantly alter zeta potential values [45]. Despite the inclusion of 5 mol% PEG-lipid in the liposome formulation, the surface charge remained negative.

Erlotinib.HCl encapsulation efficiency was determined to be 35% immediately post-manufacturing. Within the first 48 hours of storage at 4°C, a 15% loss was observed, after which the encapsulated erlotinib.HCl content stabilized. This behavior suggests the presence of two distinct populations of erlotinib.HCl within the liposomes: one fraction associated with the bilayer and another within the aqueous core. The initial loss likely corresponds to erlotinib.HCl localized in the bilayer, whereas the drug retained in the aqueous core remained stable. A similar mechanism may explain the release profile at 37°C, where a rapid release phase is observed between 0 to 4 hours, corresponding to the diffusion of bilayer-associated erlotinib.HCl. This phase is followed by a plateau, potentially reflecting erlotinib.HCl is being retained in the aqueous core, likely in a precipitated state, rendering it less susceptible to leakage. However, elevated temperatures and the acidic internal pH may contribute to increased bilayer permeability, facilitating eventual erlotinib.HCl release over time.

The hypothesis of two distinct populations of erlotinib.HCl is further supported by Cryo-TEM images, which revealed the presence of tube-shaped structures that were absent in blank liposomes. These structures are likely attributable to drug precipitation within the aqueous core and are consistent with observations reported for other liposomal formulations with remotely loaded APIs, such as the coffee bean-shaped liposomes characteristic of Doxil^®^ [46]. However, their absence in all liposomes suggests an incomplete or imperfect erlotinib.HCl loading, highlighting the need for further optimization of manufacturing parameters such as time of loading, temperature, or ammonium sulfate’s molarity.

Finally, a comprehensive biological assessment of the liposomes, especially regarding efficacy and toxicity, was essential to our study. Nanomedicine has been and continues to be a useful tool in anticancer treatment since the development of Doxil^®^. However, the frequent failure of treatments during clinical trials highlights significant challenges in clinical translation [18]. A major limitation lies in the insufficient understanding of nanomedicine interactions within the human body. Current methods for evaluating treatment efficacy rely primarily on cell lines or mouse models, which lack the complexity and physiological relevance of human tumors. Similarly, the evaluation of nanomedicine’s penetration, particularly for larger cargo systems compared to free compounds, requires validation in models of higher complexity. In this study, we utilized PDOs derived from primary ccRCC tumors, which better replicate the complex metabolic and proliferative patterns observed in human tumors. These organoids capture the native tumor architecture, with hypoxic, low-proliferative cells in the core and highly proliferative cells at the periphery, thus offering a physiologically relevant platform to evaluate the efficacy and penetration of nanomedicines.

In the 2D model, the treatment toxicity was assessed by WST-1 assay. This assay is based on mitochondrial activity and used as a surrogate to evaluate both proliferation rate and energy metabolism [47]. Erlotinib.HCl-loaded liposomes were found to inhibit non-malignant cell growth but did not affect cancerous cells. This lack of selectivity may be attributable to several factors. One explanation is the resistant nature of cancer cells compared to non-cancerous cells. Cancerous cells have genetic dysregulations that lead to a higher cell proliferation rate [48]. Furthermore, the lack of active targeting mechanisms in the liposome formulation limits its specificity. Additionally, ccRCC cell lines such as 786-O have different sensitivity to erlotinib.HCl and respond to it in a dose-dependent manner. A report showed a 60 % reduction in proliferation rate in 786-O versus only 30 % for A498 cells at a concentration of 10 µM, with the effect being primarily cytostatic rather than cytotoxic on 786-O [49].

In PDOs, although based on different principles, both WST-1 and CTG assays are routinely employed to evaluate the toxicity of compounds. After 72 hours of treatment with erlotinib.HCl-loaded liposomes, a marked reduction in viability was observed in cancerous organoids, with ATP levels decreasing to 40%. In contrast, the mitochondrial activity in 786-O cancer cell line remained above 80% in 786-O cells. This discrepancy underscores the added value of PDOs as a physiologically relevant model for assessing efficacy and toxicity. Finally, there was comparable toxicity between erlotinib.HCl-loaded liposomes and free erlotinib.HCl.

The cellular uptake of liposomes was also assessed, revealing a higher uptake in the cancerous 786-O cell line compared to the non-malignant HEK-293T cells. This result is unexpected, given the negative surface charge of the liposomes. Typically, cancer cells carry a negative surface charge, which would generally be anticipated to result in a repulsive interaction between the cells and the particles [40]. One possible explanation for the preferential uptake in cancer cells is the presence of TPGS, which has been reported to enhance the cellular uptake in several cancer cell lines such as C6-glioma cells [50], Caco-2 [51] and MCF-7 cells [52]. However, none of these studies included a direct comparison with healthy cell lines, leaving the mechanism behind the selective uptake observed here an open question necessitating further investigation.

Erlotinib.HCl’s concentration in the supernatant was measured as an indirect indicator of cellular drug uptake. The residual erlotinib.HCl concentration in the supernatant was similar between HEK-293T and 786-O cells. In 786-O, liposomal uptake was observed after 45 minutes, while a decrease in erlotinib.HCl’s concentration in the supernatant was noted after 1 hour. Although slightly delayed by 15 minutes, these results are consistent. However, the drop in erlotinib.HCl concentration at the 1-hour mark also suggests a drug uptake by HEK-293T cells, which is inconsistent with the absence of liposomal uptake observed in this cell line. This result may indicate that, upon incubation with cells, a fraction of erlotinib.HCl is released from liposomes and is taken up by cells. In the case of 786-O cells, both liposomal and free drug uptake appear to occur.

In PDOs, the liposomal uptake was notably faster and more efficient compared to 2D cell cultures. Liposomal uptake was detectable in both cancerous and non-malignant PDO models as early as 15 minutes post-incubation. Notably, the cancerous PDOs displayed higher staining intensity relative to non-malignant PDOs. This pattern suggests that, as observed in 2D models, the presence of TPGS may induce a preferential uptake by cancerous systems.

## 5. Conclusion

Erlotinib.HCl-loaded liposomes were successfully formulated. To our knowledge, this is the first instance of erlotinib.HCl being loaded actively. Results from release kinetics and Cryo-TEM support the hypothesis of erlotinib.HCl being localized at least partially as a precipitate in the aqueous core. *In vitro* studies for non-malignant systems showed toxicity in 2D cells but not in PDOs. Moreover, the cellular uptake differed significantly between 2D cell lines and PDOs in non-malignant systems. These differences underline the importance of more complex models to accurately assess particle behavior before clinical translation. More optimization of the formulation may be achieved to increase encapsulation efficiency, reduce drug release and increase cell specificity. The results provide a valuable baseline and formulation insights for the encapsulation of other TKIs containing a weak base. Such strategies could help reduce their side effects, prolong their blood circulation, and reduce the number of rounds of administration needed for therapy.

## Acknowledgments

Nirosiha Sivaruban performed experiments on erlotinib.HCl loading, helpful in building the strategy for the experiments documented in this manuscript.

